# Colonial ascidians strongly preyed upon, yet dominate the substrate in a subtropical fouling community

**DOI:** 10.1101/512699

**Authors:** Laurel Sky Hiebert, Edson A. Vieira, Gustavo M. Dias, Stefano Tiozzo, Federico D. Brown

## Abstract

Higher diversity and dominance at lower latitudes has been suggested for colonial species. We verified the latitudinal pattern in species richness of ascidians, finding that higher colonial-to-solitary species ratios occur in the tropics and subtropics. At the latitudinal region with the highest ratio, in south-eastern Brazil, we confirmed that colonial species dominate the space on artificial plates in two independent studies of five fouling communities. We manipulated settlement plates to measure effects of predation and competition on growth and survivorship of colonial vs. solitary ascidians. Eight ascidian species were subjected to a predation treatment, i.e. caged vs. exposed to predators, and a competition treatment, i.e. leaving vs. removing competitors, to assess main and interactive effects. Predation had a greater effect on growth and survivorship of colonial compared to solitary species, whereas competition did not show consistent patterns between the two life histories. We hypothesize that colonial ascidians dominate at this subtropical site despite being highly preyed upon because they regrow when partially consumed and can adjust in shape and space to grow into refuges. We contend that these means of avoiding mortality from predation can have large influences on the diversification patterns of colonial species at low latitudes, where predation intensity is greater.

## Introduction

Colonial animals reproduce clonally to construct colonies out of repeated, functionally autonomous modules, called zooids or polyps, that remain physiologically attached to one another [1]. Coloniality occurs in a wide range of animal taxa, such as corals, hydroids, bryozoans, entoprocts, pterobranch hemichordates, and ascidians [2]. While sponges are often not considered true colonies, they are modular organisms that share many of these characteristics. Modular construction and clonal growth enable escape from senescence, extreme regenerative capacity, and allow for decoupling between aggregate size and constraints on module size, potentially permitting indeterminate growth [3–5]. In some species, colonies can fuse to form chimeras [3]. Because colonial and solitary organisms differ in their reproductive and growth strategies, they differ in their abundance and diversity, depending on the environment [6]. Colonial species are generally excluded from the intertidal zone and soft substrata of marine environments, but they dominate the hard-substratum communities of many shallow seas and frequently exhibit higher diversity than solitary forms [7]. Colonial species are reported to be more abundant and diverse at lower latitudes compared to solitary species [6–9]. For example, stony corals and ascidians, which comprise both solitary and colonial forms, show a ratio of colonial to solitary species of 0.75 in temperate waters near Great Britain and 2.97 in the Caribbean [6,10]. What drives these trends in colonial vs. solitary abundance and diversity? The observed biogeographical differences in colonial and solitary forms suggest adaptations of these two life forms to selective pressures that vary with habitat and latitude.

It is widely accepted that biotic interactions such as predation, herbivory, and competition, decrease in intensity with latitude [11]. Colonial species use different strategies to survive these predatory and competitive interactions compared to solitary species. Colonial species exhibit exponential growth and can propagate indeterminately when unconstrained, giving them a competitive advantage over solitary species [2,7,12]. For example, Jamaican reef ectoprocts settle and overgrow solitary species, such as serpulids and bivalves [7]. Colonial species are also generally less susceptible to fouling and overgrowth than solitary species. For instance, colonial ascidians in the Bermuda islands were found to be less susceptible to recruitment of epibionts [13]. Colonial propagation allows for increased viability after significant damage from predation, whereas the ability of solitary species to survive damage is orders of magnitude less [6]. For example, cheilostome bryozoans in Scotland may survive after tissue damage, while nearby urchins and polychaetes show higher susceptibility [14]. Since colonial species differ from solitary species in their abilities to survive competition and predation, which both vary with latitude, this suggests that these ecological factors may underlie the trends in colonial and solitary species diversity and abundance.

Ascidians (phylum Chordata, subphylum Tunicata, class Ascidiacea) are one of the few animal groups that possess closely-related solitary and colonial species, providing an opportunity to employ a comparative approach to shed light on the ecological significance of solitary and colonial life forms. Ascidians are sessile and inhabit a range of natural (i.e. rocky shores, coral reefs) and artificial (i.e. floating docks, ship hulls) substrates worldwide [15]. They include numerous invasive species, with non-indigenous ascidians often comprising a dominant component of fouling communities in marinas [16,17]. Among the colonial taxa, ascidian species exhibit a range of forms, from stalked, upright and mound-building and multi-lobed forms to flat sheet-like species [18]. Colonial species also show differences in colony integration; some species are “social”, with individual zooids only connected at the base, while other species are highly-integrated, with zooids embedded in a common tunic. Colony forms and levels of integration are widespread. Of the three ascidian suborders, two (Stolidiopbranchs and Phleobobranchs) possess both colonial and solitary members. The third suborder, the Aplousobranchs, lack solitary forms, with the exception of one genus in which budding was lost [19]. It has been documented that colonial ascidians, among other colonial species, dominate the substrate in the tropics [7]. Colonial ascidian species also outnumber solitaries in tropical environments, characterizing 80% of species diversity [15,20]. In contrast, in most temperate systems and higher latitudes, solitaries comprise the majority of the ascidian species, though there are exceptions.

In order to further document the pattern colonial vs. solitary ascidian species richness across latitude, we analysed worldwide distribution data of ascidians. In order to understand how biotic interactions influence the dominance and richness of colonial and solitary species, we examined the ascidians of a fouling community in south-eastern Brazil, corresponding to the latitude range with the largest proportion of colonial ascidian species from the distribution analysis. While this study was conducted in artificial marinas, the diversity and access to animals at the chosen sites provides a unique opportunity to examine the ecology of solitary and colonial ascidians, which could be further tested in natural systems. We showed a general dominance of colonial animals on the substrate at this subtropical location. Lastly, we conducted a manipulative experiment to measure the effects of biotic interactions, i.e. competition and predation, on the fitness of individual solitary ascidians and colonies measured by survivorship and growth. We selected species of different growth forms and levels of colonial integration, including social species and two morphological types of highly integrated colonies in order to examine the ecological effects on growth of these different morphological strategies. Effects of predation on competition were analysed on benthic stages only, excluding any possible effects on larval stages and initial recruitment into the fouling community. By using flat panels, we also excluded effects of pre-existing spatial complexity. Our results pointed to predation as a significant selective force on colonial forms, influencing both their distribution and abundance.

## Methods

### Analysis of latitudinal distribution patterns of colonial and solitary ascidian species

Ascidian data was retrieved from the Ocean Biogeographic Information System (http://www.iobis.org). Entries were checked for valid genus names against the Ascidiacea World Database (http://www.marinespecies.org/ascidiacea/). Synonymized names were updated. Species were coded as solitary or colonial. Ratios of colonial to solitary species counts were calculated for each 10° latitudinal bin (Figure 1).

**Figure 1:**
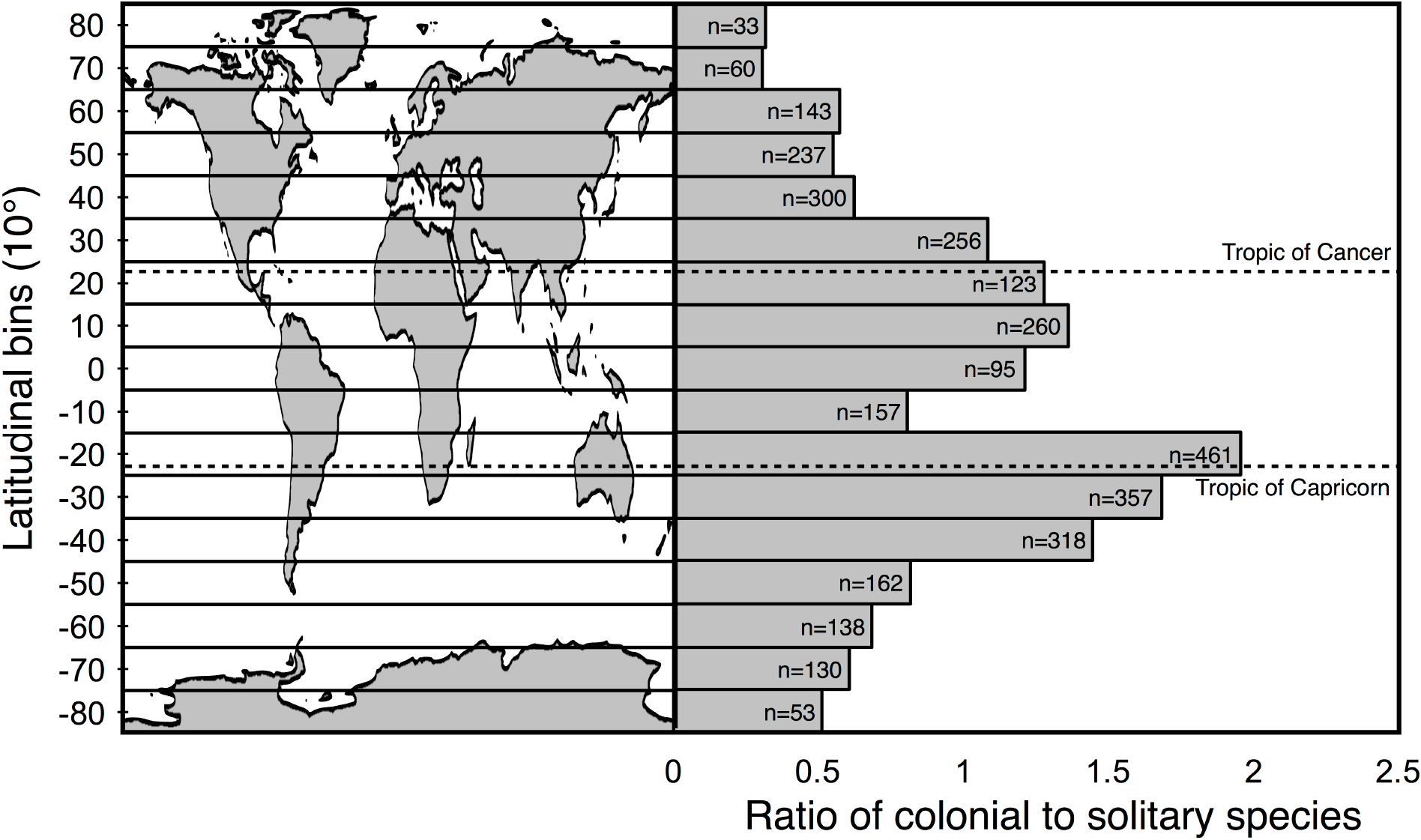
Ratio of colonial to solitary ascidian species counts within 10° latitudinal bins. Based on data from the OBIS (Ocean Biogeographic Information System) database. Number of total ascidian species (n) in each bin is indicated on the bars.

### Determination of dominance and species richness of colonial vs. solitary species in a SE Brazil fouling community (2007-2010 field studies)

We examined the coverage and species richness of sessile colonial and solitary species of all animal phyla at a subtropical latitude where the proportion of colonial to solitary ascidian species was the highest (in the 15-25° S latitudinal bin), with almost twice the species of colonial compared to solitary ascidian species. To determine patterns of dominance and richness of colonial vs. solitary forms in the São Sebastião Channel, we included in this study data obtained from plates exposed to predators at five distinct sites from the years 2007 (Segredo) and 2010 (Feiticeira, Figueira, Yacht Club Ilhabela-YCI and Curral) (Figure 2). At all sites, panels were suspended (at least 1.5 m deep and 2 m from the bottom) from artificial structures (moorings at Segredo; jetties at Curral, Feiticeira and Figueira; and floating platforms at YCI), which are covered by a diverse fouling community that can supply settlers to the experimental plates. From 30 × 30 cm plates in Segredo, and 15 × 15 cm plates in Curral, Feiticeira, Figueira and YCI, we used the 10 × 10 cm central area to register the number of species and to take photos to calculate the percent cover of colonial and solitary taxa present (for taxa list, see supplementary materials). For Segredo (2007), data were obtained from the same plates in different sampling events, and therefore coverage and richness were compared between life forms (colonial × solitary) and across time (30, 60, 90, 120, 150 and 180 days; n = 6) using a Repeated Measures ANOVA. Repeated measures ANOVA was conducted using R [21] with the ezANOVA package [22]. Mauchly’s Test of Sphericity indicated that the assumption of sphericity had been violated for the coverage dataset, (coverage: W= 0.002, *p* < 0.05; richness: W= 0.182, *p* < 0.05) thus Greenhouse-Geisser corrections were applied (coverage: ε = 0.369, richness: ε = 0.621). [23,24]. For significant interactions we highlighted the differences in a descriptive way [25]. 2010 data were obtained from independent sets of panels after 30 (n = 12) and 100 days (n = 18), with coverage and richness being compared between life forms and across time for Feiticeira, Figueira, and YCI, and only between life forms for Curral (all panels were lost after 30 days), using a two- or one-way ANOVA in the program Systat. In most analyses data normality and homoscedasticity was not achieved even after square-root transformation, but, as ANOVA tests are robust for such deviations when data are balanced and sufficiently replicated [26], we decided to use an ANOVA because it is more powerful compared to non-parametric tests [25].

**Figure 2:**
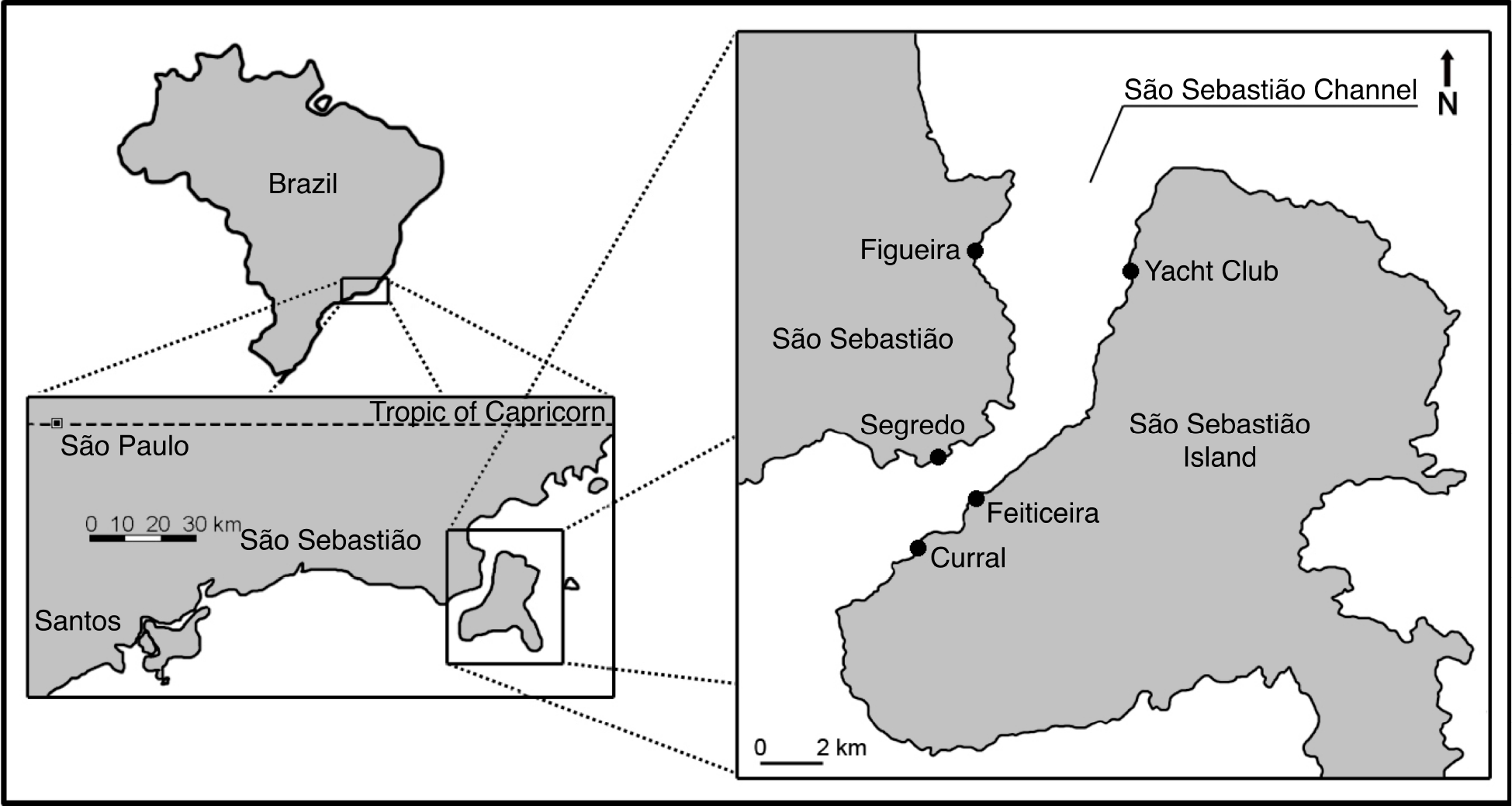
Map of SE Brazil Study Locations. Experimental panels were deployed at Yacht Club docks (23°46'29''S, 45°21'25''W), piers at Feiticeria (23°50'43''S, 45°24'40''W), Curral (23°52'2''S, 45°26'2''W), and Figueira (23°53'3'' S, 45°16'48''W), and an artificial mooring at Segredo (23°49'44''S, 45°25'24''W).

### Testing effects of predation and competition on growth and survivorship of colonial and solitary ascidians (2016 field studies)

#### (a) Species selection

We selected representative species of the ascidian fouling community along the São Sebastião Channel because of their abundance and to capture as much diversity of life histories and forms as possible to limit results that may be due to morphological or phylogenetic constraints (Figure 3). Species were selected from five ascidian families (Styelidae, Pyuridae, Clavelinidae, Polyclinidae and Didemnidae) belonging to two different orders (Stolidobranchia and Aplousobranchia). Solitary species include species belonging to two different families: *Styela plicata* and *Herdmania pallida*. The six colonial species represent different degrees of zooid integration/separation; four species bear zooids arranged into systems with shared cloacal aperture and embedded completely in a common tunic (*Polyclinum constellatum, Aplidium accarense Didemnum galacteum* and *Botrylloides niger*). Of those, one has a colony-side vascular system, *Botrylloides niger*, and thus represents a high level of colonial integration since zooids are physiologically interdependent. Each species can also be categorized by “morphological strategy” or “functional group” based on colony morphology [6,27]. Two species connected by basal tunic or stolons only at their bases (*Clavelina oblonga* and *Polyandrocarpa zorritensis*), and grow as “trees” or “runners” since they grow as either branching encrusted or erect, and occasionally the branches become so dense that they form more massive “mounds”. The other species grow either as “sheets”, encrusting the substrate in two dimensions, or as “mounds” when they possess a significant vertical dimension to their growth (Figure 3). Whereas the biota present at this highly diverse subtropical site did not provide replicate pairs of evolutionary divergent species to avoid lineage specific effects that may be unrelated to form, it did provide species that are distributed among major monophyletic groups of ascidians: the two solitary species used are in the families Styelidae and Pyuridae, the two mound-like colonial forms are both aplousobranchs in the family Polyclinidae, whereas the two social and two sheet-like compound forms are divided between aplousobranchs and stolidobranchs (Figure 3) [18,28]. We used only adults of the different life forms to remove effects of factors like larval supply and post-settlement mortality.

**Figure 3:**
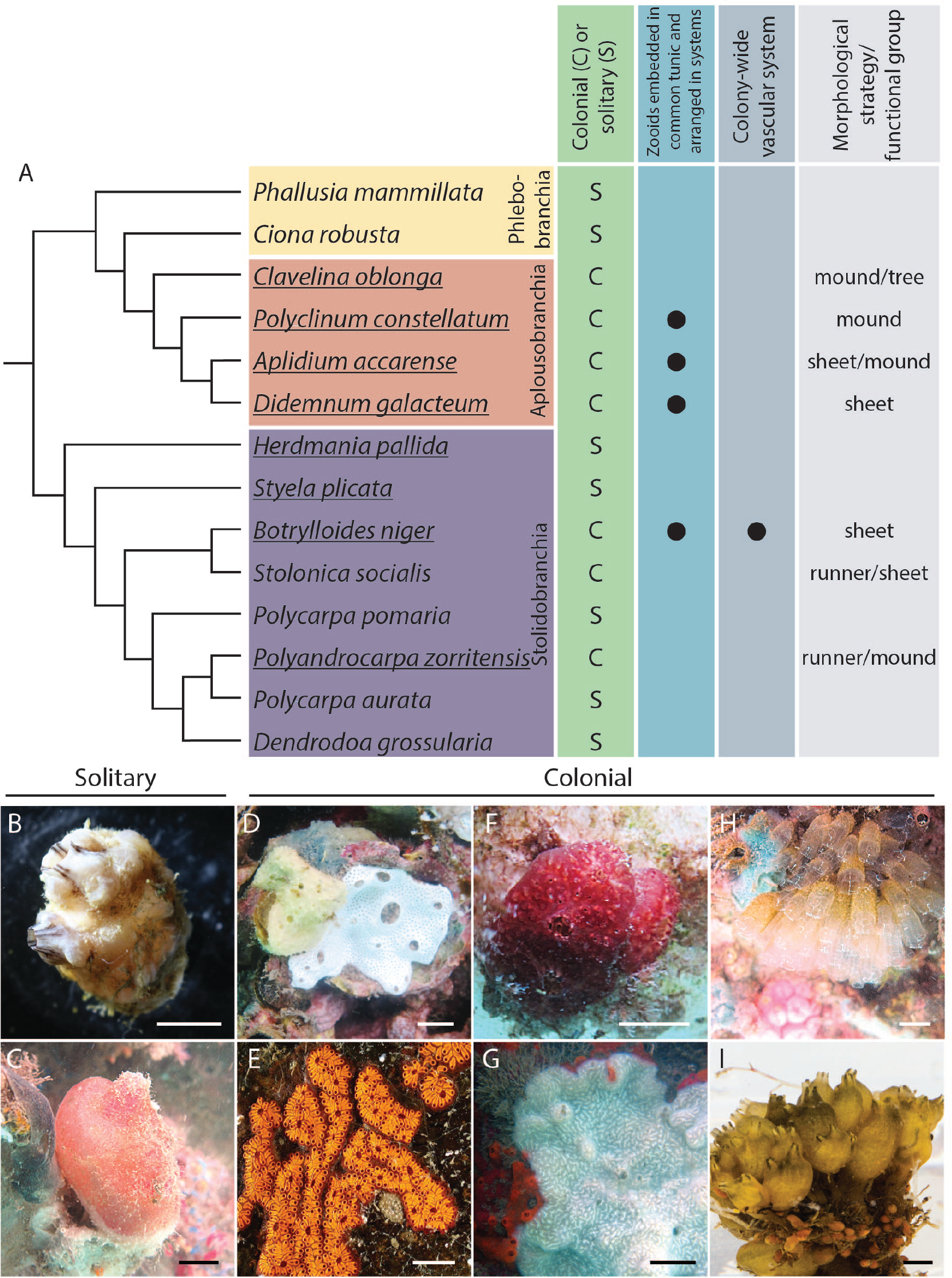
Phylogeny and photos of ascidian species used in study. Simplified phylogeny of the ascidians based on Delsuc et al. (2017) and Alié et al. (2018) [65,66]. Underlined species are those used in this study. Boxes around groups of species are labelled to demarcate ascidian orders. Column to the right of the phylogeny indicates whether each species is colonial (labelled with a “C”) or solitary (labelled with a “S”). Additional columns indicate degree of colony integration by listing if zooids are embedded in a common tunic and arranged into systems around a shared cloacal opening, whether colonies possesses shared vasculature, and which morphological strategy/functional group category each species falls into. (B-J) Photos of experimental species. (B-C) Solitary species. (B) *Styela plicata.* (C) *Herdmania pallida.* (D-E) Colonial - sheet species. (D) *Didemnum galacteum*. (E) *Botrylloides niger.* (F-G) Colonial - mound species. (F) *Polyclinum constellatum.* (G) *Aplidium accarense*. (H-I) Colonial - social species. (H) *Clavelina oblonga*. (I) *Polyandrocarpa zorritensis*. Scale bars: (B-C) approx. 10 mm, (E) 14 mm, (G) 30 mm, (I) approx. 5 mm.

#### (b) Collection and culturing for manipulative field experiment

Solitary and colonial species were collected and cultured for three months in preparation for the manipulative experiment described below at the YCI (February - May of 2016; Figure 2). For solitary species, gametes were dissected from a single pair of reproductive adults and in vitro fertilization was conducted [29]. Larvae were allowed to settle on petri dishes, which had previously been scraped with sandpaper to provide a rough settlement surface. The petri dishes containing the metamorphosed juveniles were then attached onto temporary PVC panels and were allowed to grow at Praia do Segredo for three months in cages submerged at 2 m. Three-month-old juveniles were used to begin the manipulative experiment described below. For colonial species, small pieces of colonies were initially “planted” to the centre of the underside of a temporary PVC panel with plastic cable ties and allowed to grow into a larger colony for three weeks. Large colonies were sub-cloned into smaller colonies (i.e. cut into smaller genetically-identical pieces) of approximately 1.5 × 1.5 cm, which were “re-planted” onto the final PVC panels used for the manipulative experiments as described below.

#### (c) Experimental design and data collection

The experiment was conducted along a single stretch of floating concrete platforms at the Yacht Club of Ilhabela from May to August, 2016. Each species was attached to a 30 × 30 cm PVC panel and suspended horizontally 3 m deep and 1 m apart (see Figure 4). For solitary species, four or six three-month-old juveniles of *S. plicata* or *H. pallida* respectively were attached 5 cm apart with superglue to the downward face of the same panel (Figure 4E). For the two social species (*C. oblonga* and *P. zorritensis*) five zooids connected by their tunic were attached to the centre of the panels using two small cable ties (Figure 4D). For the other colonial species, a single large colony was sub-cloned into 1.5 × 1.5 cm squares and “planted” to the centre of the panels as described above (Figure 4C). Predation was manipulated using screen cages (30 × 30 × 8 cm, and 0.75 × 0.75 cm mesh) (Figure 4B). Half of the panels were enclosed by full-cages, excluding large predators (mainly fishes), while the other half were enclosed by open-cages, which was the same of full-cages but lacking the roof, allowing then the access of predators while controlling for any eventual cage artefact [30] (Figure 4).. Competition was manipulated by removing all surrounding settlers using a paint scraper every two weeks, since fouling species settled but did not have time to grow significantly into the focal specimen during this time window. Competitors included barnacles, mussels, bryozoans, and other ascidians. Half of the panels were cleaned of these competitors and the remaining half were left uncleaned. The panels were randomly distributed along the floating platforms. For each species, predation and competition treatments were varied orthogonally to assess main effects and interactions. For each of the eight species, four replicates were allocated to four treatments that combined predation and competition: “full-cage/cleaned,” “full-cage/uncleaned,” “open-cage/cleaned,” and “open-cage/uncleaned” (Figure 4).

**Figure 4:**
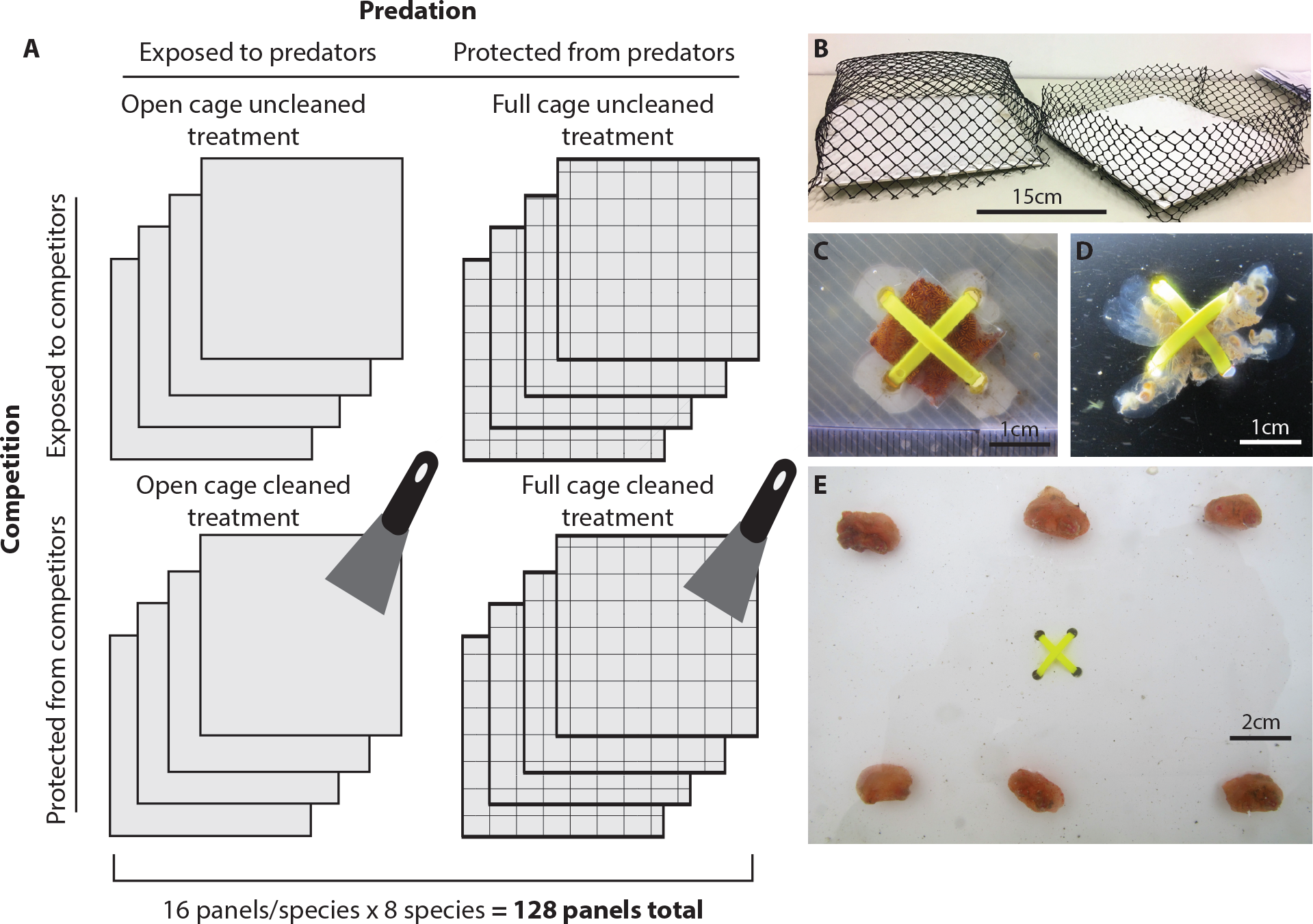
Design of experiment to manipulate predation and competition. (A) Schematic of experimental panels in predation and competition treatments showing 4 replicate panels per treatment. (B) Photo showing closed cage panels (left) and open cage panels (right). (C) Example to indicate how pieces of colony are attached to centre of panel by two cable ties. (D) Example to indicate how a cluster of connected zooids is attached to the centre of a panel with cable ties. (E). Example to indicate how juvenile solitary species were attached to a panel.

Survivorship of each focal species was recorded, and photographs of the panels (with ruler) were obtained every other week. At the same time, the panels in the “cleaned” treatments were scraped and the cages were cleaned to prevent build-up of fouling species. The area of the focal species (i.e. the region of interest) was measured in the photos using ImageJ (Wayne Rasband, National Institutes of Health, Bethesda, MD, USA). In order to assess whether the area measurements were reliably capturing the growth of the ascidians, we determined if area was correlated to dry mass of each species. For dry mass measurements, samples were baked in an oven at 60°C for 24 hours. At six weeks, for each solitary species, one individual from each panel was removed for dry mass analyses. At the end of the experiment, all remaining specimens, including both solitary and colonial species, were collected and their dry mass was determined. Using these data, we generated area-by-dry-mass growth curves. Plots show that area is a correlate of mass for most species (see electronic supplementary materials, Figure S1). For one species, *P. zorritensis*, there were too few data points to generate a correlation.

#### (d) Statistical analysis

Only the first eight weeks of data were used in our analyses because growth rates peaked during this time and storms during the last month damaged many panels and reduced the replication. Differences between treatments were already clear in just the first 8 weeks of the experiment.

We tested if survivorship of the two different life forms was associated with the combination of predation and competition treatments using a Chi-square approach. For each life form and time (2, 4, 6, and 8 weeks), we separately obtained the expected survivorship by averaging the survivorship from the four treatment combinations and compared it to the observed values. Since the data were not independent in time for each life form, the critical *p*-value was corrected using the Bonferroni adjustment for four different tests, with a corrected value of *p* = 0.0125 for a test using a Type I error of 5% [25].

In order to compare the effects on growth of each treatment among life forms, the area of each species at each time point was standardized by converting the raw “region of interest” into z-scores. Pseudoreplicates of the solitary forms were removed from the data and only one individual per panel was randomly selected and used in the analyses. Repeated measures ANOVA was conducted as before. Mauchly’s Test of Sphericity indicated that the assumption of sphericity had been violated, W= 0.3398, *p* < 0.05, thus Greenhouse-Geisser corrections were applied (ε = 0.6585).

## Results

### Latitudinal trends in colonial and solitary ascidian species counts

The highest proportion of colonial to solitary ascidian species was 1.96 in the 15-25° S latitude bin (Figure 1). The lowest proportion was 0.30 colonial to solitary species in the 65-75° N latitude bin. Solitary species outnumbered colonial species (ratio < 1) in the 45-55° N bin and further north and in the 35-45° S bin and further south. The tropical and subtropical regions generally showed a higher proportion of colonial species compared to solitary species (ratio >1), with the exception of the 5° N-5° S latitudinal bin, in which solitary species outnumbered colonial species. However, the colonial-to-solitary species ratio was still higher in the 5° N-5° S bin than in most temperate and polar regions. The general trend was a gradient with decreasing colonial-to-solitary species ratios toward the poles, but two peaks on either side of the equator.

### Dominance and richness of colonial and solitary species in south-eastern Brazil

Two of us (EAV & GMD) have been studying community structure on settlement plates in the São Sebastião Channel in the state of São Paulo (SP) in south-eastern Brazil for over a decade [30–33]. In the present study, we documented that colonial animals dominate over solitary forms and show higher species richness at five sites in two independent investigations in the SE Brazil fouling community (Figure 5, Supplementary Tables 1 and 2, data in supplementary materials). These separate investigations took place in two separate years and on panels in which animals were exposed to predators. In 2007 at Segredo, colonial species reached 70-90% cover of panels by 60 days and continued at this coverage while solitary species remained near 10-20% cover. Only in the first 30 days of submergence did the solitary species exhibit high coverage, but colonial forms quickly overtook them in the next month and throughout the experiment. Although showing the same number of colonial and solitary species in the first 30 days, colonial species outnumbered solitary species at Segredo after 60 days, being almost 6 times more numerous after 180 days (Figure 5A, Supplementary Table 1). In 2010 at four additional sites, Feiticeira, Curral and Figueira piers, and YCI floating docks, colonial species displayed higher percent cover and richness compared to solitary species at both 30 and 100 days of submergence (Figure 5B, Supplementary Table 2). All sites showed less than 30% cover of solitary species (except for panels at 30 days of submergence at Segredo), while the majority of sites across time points showed over 50% cover of colonial species. The trends in the effects of submergence time on solitary vs. colonial coverage varied among sites. At Segredo colonial coverage and richness increased between 30 to 90 days, while solitary species decreased with submergence time; Feiticeira and YCI showed an increase in coverage and number of both life forms between 30 and 100 days; Figueira showed an increase of both colonial and solitary species number but no changes in cover between 20 and 100 days. Despite some variation, these results demonstrate a general dominance and diversity of colonial life histories during the establishment of fouling communities that is maintained over the years at a subtropical site.

**Figure 5:**
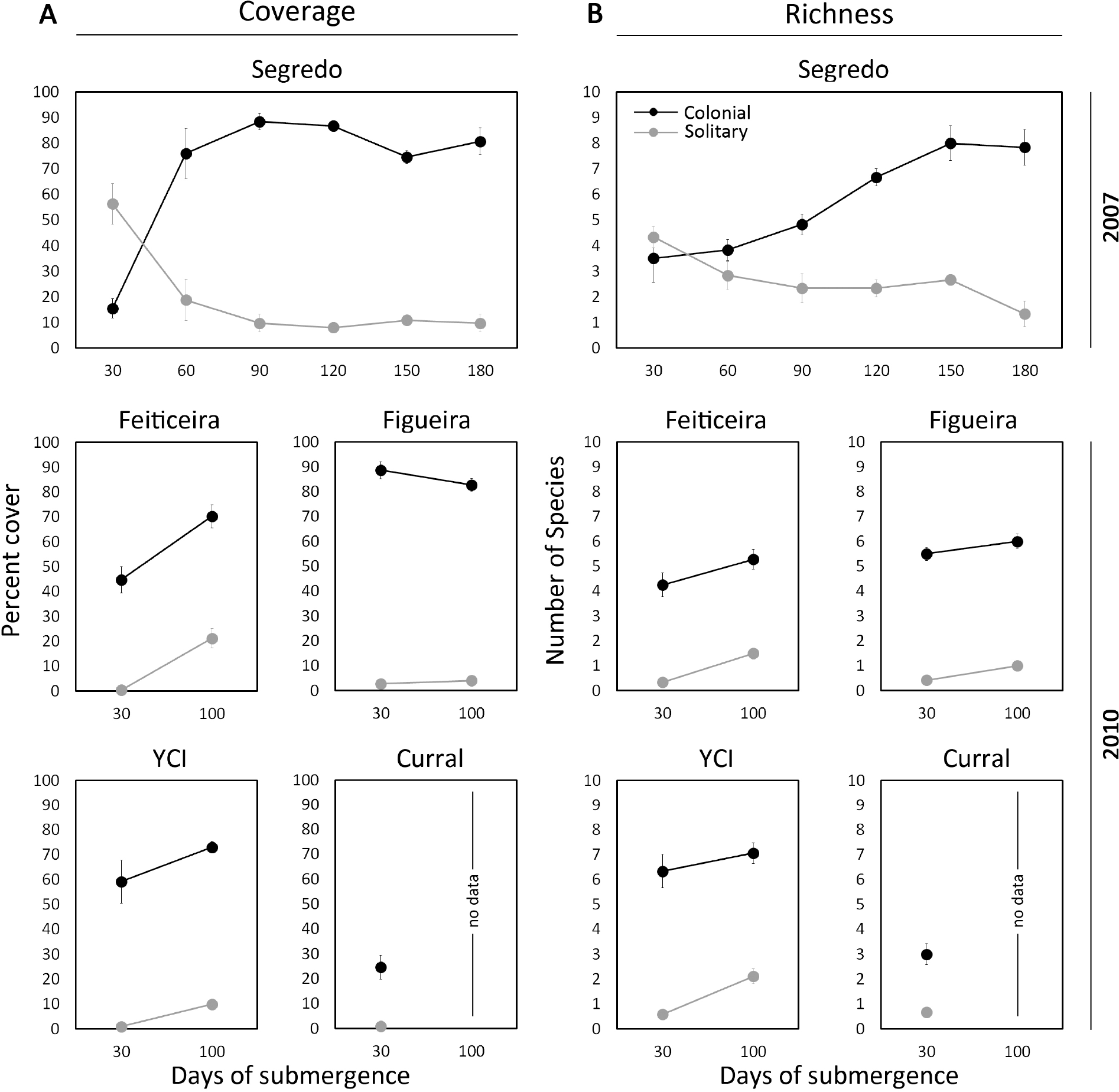
Colonial forms dominate the substrate in the SE Brazilian fouling community. (A) Mean percent cover (± SE) and (B) mean richness (± SE) of colonial (black) and solitary (grey) fouling species in panels over 180 days of submergence at Segredo in 2007 and after 30 and 100 days of submergence at Feiticiera, Figueira, YCI, and Curral in 2010.

### Effects of predation and competition on colonial and solitary ascidians

Overall, colonial ascidian survival was higher on predator-protected treatments throughout the study with departures from the expected average survivorship for colonial ascidians (*p* < 0.0125 for all analysed times) but not for solitary ones (*p* > 0.0125 for all analysed times) (Figure 6, data in supplementary materials). More than 60% of colonies survived after 12 weeks when protected against predation, but only 40% when exposed to predators (n=16 per treatment, Figure 6A). Around 40% or less of solitary ascidians survived the 12-week study, without a major difference in survivorship across any of the treatments (Figure 6B). Competition did not have an effect on survivorship of either life form.

**Figure 6:**
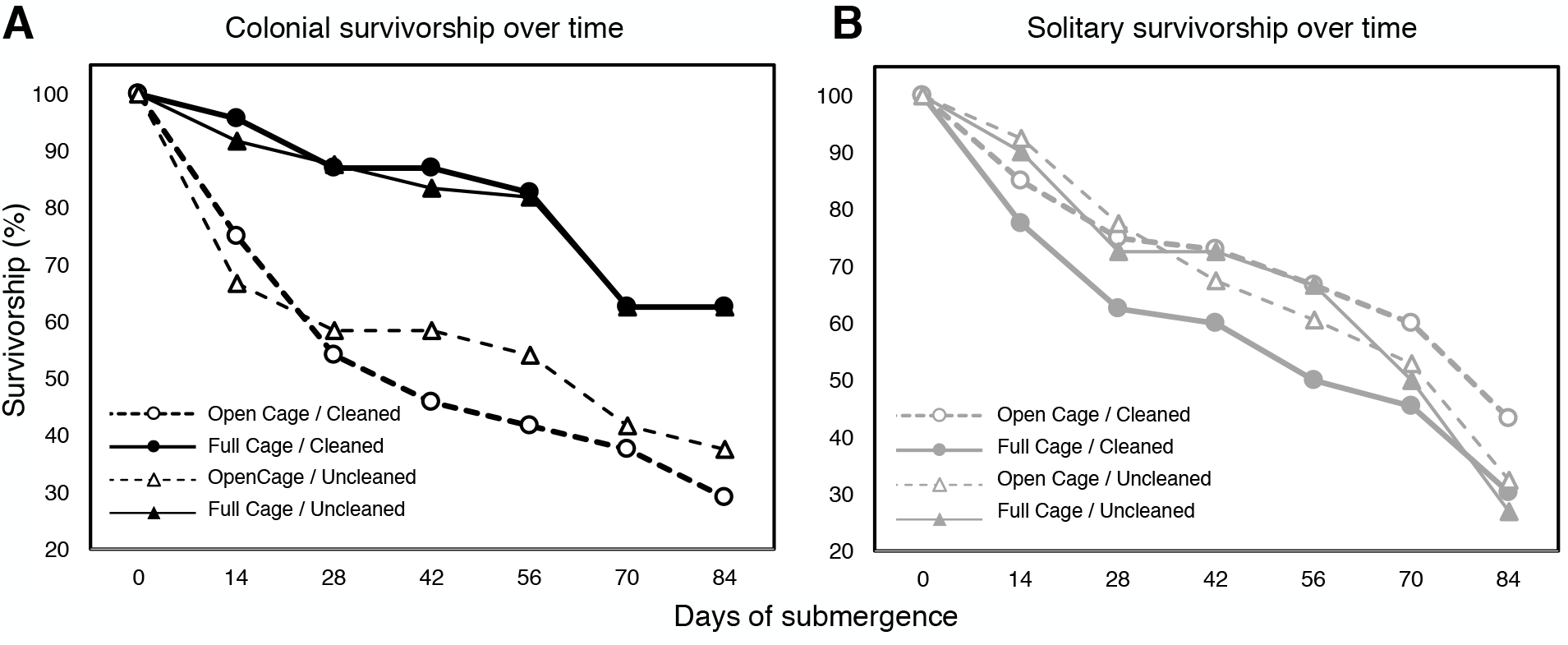
Survivorship of colonial and solitary ascidians in predation and competition treatments. (A) Proportion of colonial ascidians alive in each treatment over the 12-week study period (n=16 plates per treatment). (B) Proportion of solitary ascidians alive in same treatments in the same 12 weeks.

Colonial ascidians protected against predators achieved larger sizes than those exposed, suggesting an effect of predation on growth (Figure 7, RM-ANOVA: Form*Predation*Time, *p* < 0.05, Supplementary Table 3). Representatives of all the colonial growth forms, including mound-like (*Aplidium accarense*, *Polyclinum constellatum*), social (*Polyandrocarpa zorritensis*), and sheet-like (*Didemnum galacteum*), grew substantially more in the predator-protected treatments (Figure 7A-F). However, *Clavelina oblonga* (social) and *Botrylloides niger* (sheet-like) did not show differences between predator-protected and exposed as great as other species. Both solitary species in predator-protected treatments showed similar growth to those in the exposed treatments (Figure 7G-H). This suggests that different forms (colonial vs. solitary) respond differently to predation, with many colonials being more susceptible than solitary species, and this effect changes over time (RM-ANOVA: Form*Predation*Time, *p* < 0.05, Supplementary Table 3). Competition did not have a significant effect on growth of any life form (RM-ANOVA, Supplementary Table 3).

**Figure 7:**
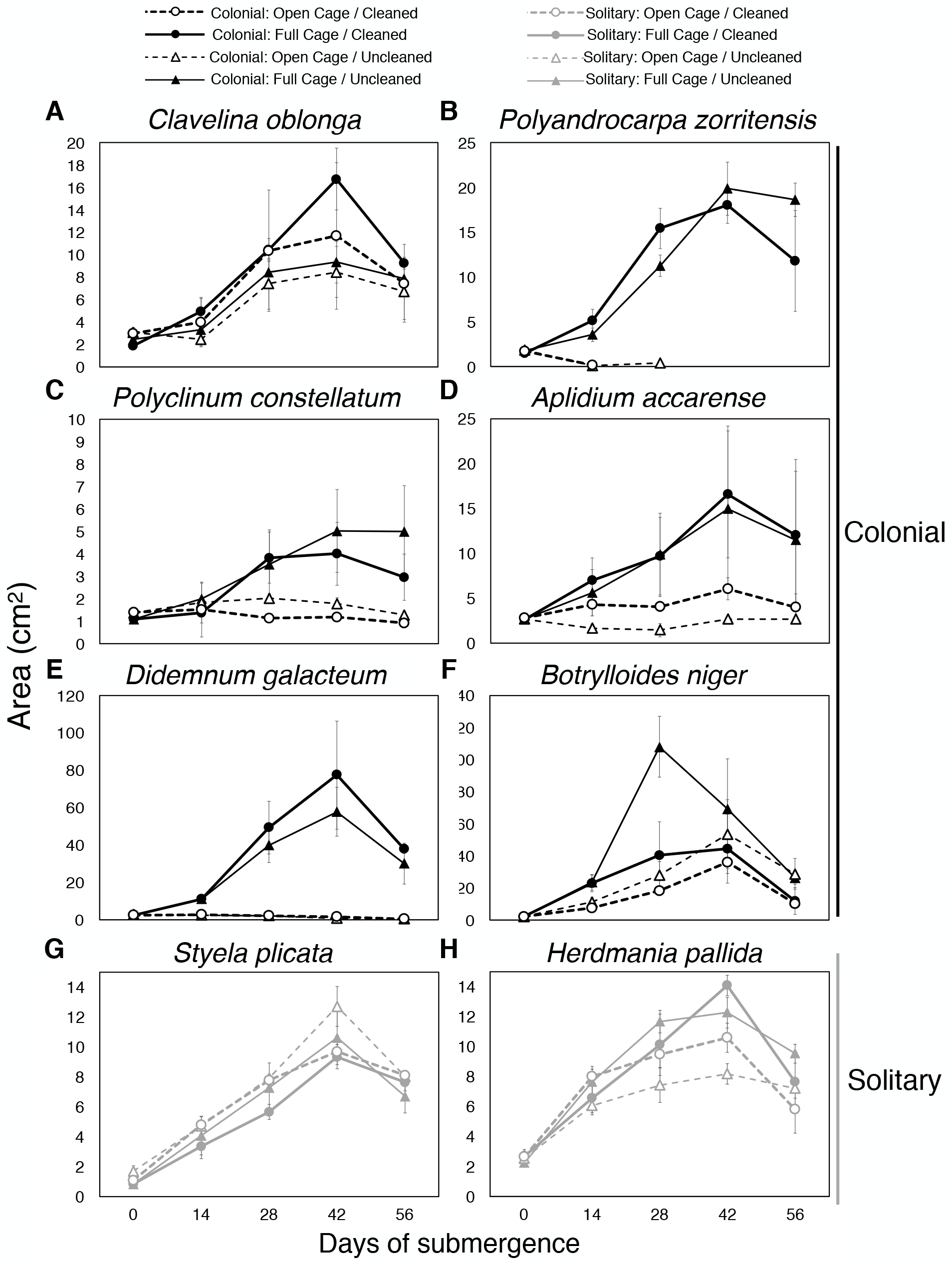
Growth of ascidian species in predation and competition treatments. Mean area (± SE) of social colonial (A-B), mound-forming colonial (C-D), sheet-like colonial (E-F), and solitary (G-H) ascidians over 56 days: (A) *Clavelina oblonga*. (B) *Polyandrocarpa zorritensis.* (C) *Polyclinum constellatum.* (D) *Aplidium accarense*. (E) *Didemnum galacteum*. (F) *Botrylloides niger.* (G) *Styela plicata*. (H) *Herdmania pallida*. As in previous figures, black lines are used for colonial species and grey are used for solitary. N=4 replicate plates per treatment

## Discussion

### Latitudinal gradient in the ratio of colonial to solitary ascidian species

It has been suggested that diversity and dominance of colonial animals are higher in the tropics (i.e. at lower latitudes) compared to solitary species, which are more abundant in temperate waters (i.e. higher latitudes), however this trend has not been clearly documented [6–9]. Colonial ascidians, in particular, have been noted to be more species rich in the tropics, comprising 80% of total species diversity [15]. Here, using OBIS data (Figure 1), we further characterized the latitudinal trends in ascidian species richness. We found that colonial ascidians are more species rich than solitary species in tropical and subtropical latitudes, confirming the pattern noted in other studies. We also found a general gradient in the ratio of colonial-to-solitary species that decreases with latitude from less than half of all ascidian species being colonial near the poles to nearly two times more species being colonial at low latitudes. However, there appears to be two peaks in the ratio - one on either side of the equator close to the subtropics. The 5° N - 5° S latitudinal bin that spans the equator, on the other hand, shows lower ratios of colonial-to-solitary ascidian species. It is unclear why the tropics show a depression in the ratio, but it may be due to some bias such as undersampling, since the number of total species recorded in this equatorial zone is slightly lower compared to nearby subtropical latitudes. Alternatively, the bimodal distribution may represent a real phenomenon, as other studies have found one or multiple peaks in marine species diversity that are offset from the equator [34]. Nonetheless, while the equatorial latitudes show a drop in the number of colonial-to-solitary ascidian species compared to surrounding latitudes, the ratio is still larger than most temperate and polar latitudes. Biotic interactions, including predation, are also known to be greater in lower latitudes [11,35,36]. Thus, we hypothesize that colonial vs. solitary species diversification may be linked to one or more ecological differences that vary with latitude.

### Colonial forms dominate and show higher species richness compared to solitary forms, yet are more susceptible to predation in a subtropical fouling community

We chose the latitude with the largest ratio of colonial-to-solitary ascidian species (i.e. subtropical SE Brazil) to examine both the dominance and richness of colonial vs. solitary species, and the effects of biotic interactions on growth and survivorship of these two life forms. Our data (restricted to artificial substrates) showed that colonial species were more diverse and dominant over solitary species in the SE Brazilian fouling community (composed mostly of non-native, introduced ascidian species), being several times more numerous and taking up to 70-90% of the substrate space. Here, we examined all species (not only ascidians) in the fouling community, since the community is made up of a number of colonial and solitary species from a range of phyla. This trend, with colonial species dominating the substrate, was consistent over multiple years and sites 2-15 km apart in SE Brazil. Although our study was restricted to artificial substrate and the resemblance with natural systems is still a matter for further investigation, the results obtained here are clear and may not only provide insights for coloniality patterns but also for invasiveness strategies, as most of the organisms growing in these artificial substrates are non-indigenous species.

By manipulating predation and competition at the same site in SE Brazil, we were able to determine if these ecological factors affected the growth and survivorship of colonial and solitary ascidians differently. Since competition has been reported to favour colonial over solitary forms [6,7], we predicted that competition would affect growth and survivorship of solitary more than colonial species. Surprisingly, our data did not show any effect of competition on either solitary or colonial life forms during the eight-weeks for which data were analysed, or any noticeable trends for the 12 weeks of the experiment. Although we have a procedural limitation of not using true settlers, and thus not allowing competition effects to take place on early and more susceptible stages, our experimental panels were fully occupied by 6-8 weeks which clearly impose some competition pressure even for larger organisms. It is possible that only heavy crowding generates a significant effect of competition on growth in our system. Therefore, our study may not have included enough time to allow sufficient crowding to capture competition effects. The only colonial ascidian species in this community for which competition effects have been documented is *Didemnum perlucidum*, but effect of competition on size is only evident after 18 weeks [37].

Our data show that predation had a greater effect on growth and survivorship of colonial compared to solitary ascidian species. Thus, colonial ascidians may employ fewer defensive strategies against predation (morphological or chemical defenses) than solitary species. This is surprising, given that ascidians are studied for the presence of defensive metabolites [13,38–41]. Few studies have directly compared the relative strength of defenses between solitary and colonial species. While colonial ascidians from Bermuda show less palatability than solitary species [13], colonial ascidians from the Gulf of Mexico have been shown to have higher palatability to fishes than solitary ascidians [42]. At our site the most observed predators are fish, including *Abudefduf saxatilis* (Linnaeus, 1758), *Stephanolepis hispidus* (Linnaeus, 1766), and *Diplodus argenteus* (Valenciennes, 1830) [30,43,44]. Here solitary species were less predated by fish likely due to anatomical defenses, such as tougher tunics or camouflage [13,42]. In addition to morphological defences, solitary species have been known to protect themselves from predation via an “escape in size” strategy, in which animals prevent predators from being able to bite them by growing rapidly past a size-sensitive threshold [7,45]. One of the solitary species used in this study, *Herdmania pallida*, has numerous tunic spicules that probably deter predators similar to other *Herdmania* species [46,47].

Solitary species may be particularly vulnerable to predation at early life stages [48–50]. For example, while 1-3-day-old juveniles (0.5−1 mm in diameter) were killed by predators (>90%) in less than an hour, 4 week young adults of the solitary ascidian *Molgula manhattensis* (approx. 5 mm) survived after exposure to predators (>90%) [50]. Mortality of early life stages of solitary ascidians was not explored in our study because the solitary species used were already three months old when the experimental regimes started.

We found that colonial species are less resistant to predators than solitary species in a region where colonial species are both dominant and more species rich compared to solitary species. We hypothesize that predation acts as a selective force on solitary forms, with only those species that can withstand predation being maintained, while predation seems to be less important in restricting the occurrence of colonial species. This may help explain why even with high predation the diversity of colonials is much higher than that of solitary species. Further, this suggests that colonial species have alternative strategies to survive predation to maintain higher dominance and diversity in this system. Colonial species, due to their unique growth mode, have extreme re-growth potential; they have the ability to survive partial or even major damage to the colony by regenerating lost parts. Some colonial species even have the potential to fuse to become chimeras, which may be considered a survival strategy since mortality often decreases with colony size [2]. Colonial species can also grow laterally to take advantages of cracks, crevices, caves, and other cryptic environments that may act as spatial refuges from predators [51,52]. Such a protective mechanism of “running away” into refuges can been termed “escape in space” [53,54]. In addition, colonial species can scale back (e.g. some undergo hibernation) when conditions are poor, then regrow [55–58]. This is similar to the idea of “temporal escape” or “escape in time” proposed by Lubchenco and Gaines (1981), in which colonies may withstand occasional disturbance, such as a dearth of resources [51] or the presence of predators, by temporally modulating growth [53,54]. As these alternative protective strategies are linked to the colonial life style, we hypothesize that coloniality provides non-defensive mechanisms to tolerate disturbances, allowing colonial species to avoid mortality and eventually dominate the substrate. In regions where predation is intense, coloniality may provide alternative protective strategies, which could explain the large diversity and dominance of these organisms even in the study area, where predation appears to be strong [30,43,59].

Defenses are known to be more common in tropical benthic ecosystems (although there are exceptions, see [60]), where it has been suggested that high rates of predation and herbivory favor more strongly defended prey than in temperate and polar regions [61]. For example, physical and behavioral defenses due to increased fish predation are known at low latitudes; toxicity of sponges and sea cucumbers is inversely related to latitude [62], littoral gastropods show reduced foraging time at lower latitudes as a means of avoidance of shell-crushing fish, which are more abundant in the tropics [63,64]. As a result of this study, we suggest that coloniality-linked survival strategies promoted by higher predation at lower latitudes (i.e. propagative regeneration or clonal expansion to escape and avoid predators) may act as drivers of colonial diversity observed in the tropics. Since predation is more intense in the tropics where diversity and dominance of colonial species is greatest, it is possible that predation is driving the origin and diversification of coloniality as a survival strategy.

### Conclusions

We first document a latitudinal gradient in the number of colonial to number of solitary ascidian species, with the highest ratios in lower latitudes. We find that in a subtropical fouling community corresponding to the latitude with the highest colonial-to-solitary ascidian ratio, colonial species are more diverse, dominate space, and predation affects growth and survival of colonial ascidians more than solitary species. We hypothesize that while solitary species rely on strategies to avoid being eaten or damaged, colonial species rely on strategies to tolerate partial injury. In particular, while solitary species avoid being eaten while in place, colonial forms rely on their abilities to regrow when partially damaged or when conditions are unfavourable and to grow into spatial refuges. Thus, coloniality provides unique protective strategies, allowing colonial species to require fewer defenses to avoid mortality and eventually dominate the substrate. Since colonial species are more specious in the tropics, where predation pressure is high, we hypothesize that predation may be an important driving force for the diversification of colonial forms.

## Supporting information

Electronic Supplementary Material, Dataset S1

Electronic Supplementary Material, Figure S1

## Data accessibility

The datasets supporting this article have been uploaded as part of electronic supplementary material.

## Authors’ contributions

L.S.H. designed and performed the field and laboratory work for the 2016 manipulative study, and performed statistical analyses for the 2016 study. E.A.V. carried out field work and analysis for the 2007-2010 field studies. G.M.D. helped with experimental design and statistical analysis. F.B. was involved in the experimental design and helped with logistics. L.S.H. wrote the manuscript with contributions from all the other authors.

## Acknowledgements

We thank Denisse Galarza, Stefania Gutierrez, Juan Jiménez, and David Soares for help with fieldwork. We are grateful to the staff of the Yacht Club of Ilhabela for allowing us to use their facilities. We also thank the technicians and staff of the Centro de Biologia Marinha da Universidade de São Paulo (CEBIMar-USP) and Richard Strathmann for providing helpful feedback on the manuscript.

## Electronic Supplementary Material

### Electronic Supplementary Material, Figure S1: Relationship between ascidian dry weight and area

Lines of best fit and R^2^ values indicating strength of relationship between dry mass and area of: (A) *Clavelina oblonga*. (B) *Polyandrocarpa zorritensis.* (C) *Polyclinum constellatum.* (D) *Aplidium accarense*. (E) *Didemnum galacteum*. (F) *Botrylloides niger.* (G) *Styela plicata*. (H) *Herdmania pallida.*

### Electronic Supplementary Material, Dataset S1: Species and growth data for all experiments

Spreadsheet containing species counts and coverage data for 2007 and 2010 experiments and growth data for 2016 manipulative experiment.

