## Supplementary figures and images for "Colonial ascidians strongly preyed upon, yet dominate the substrate in a subtropical fouling community"

### Electronic Supplementary Material, Figure S1

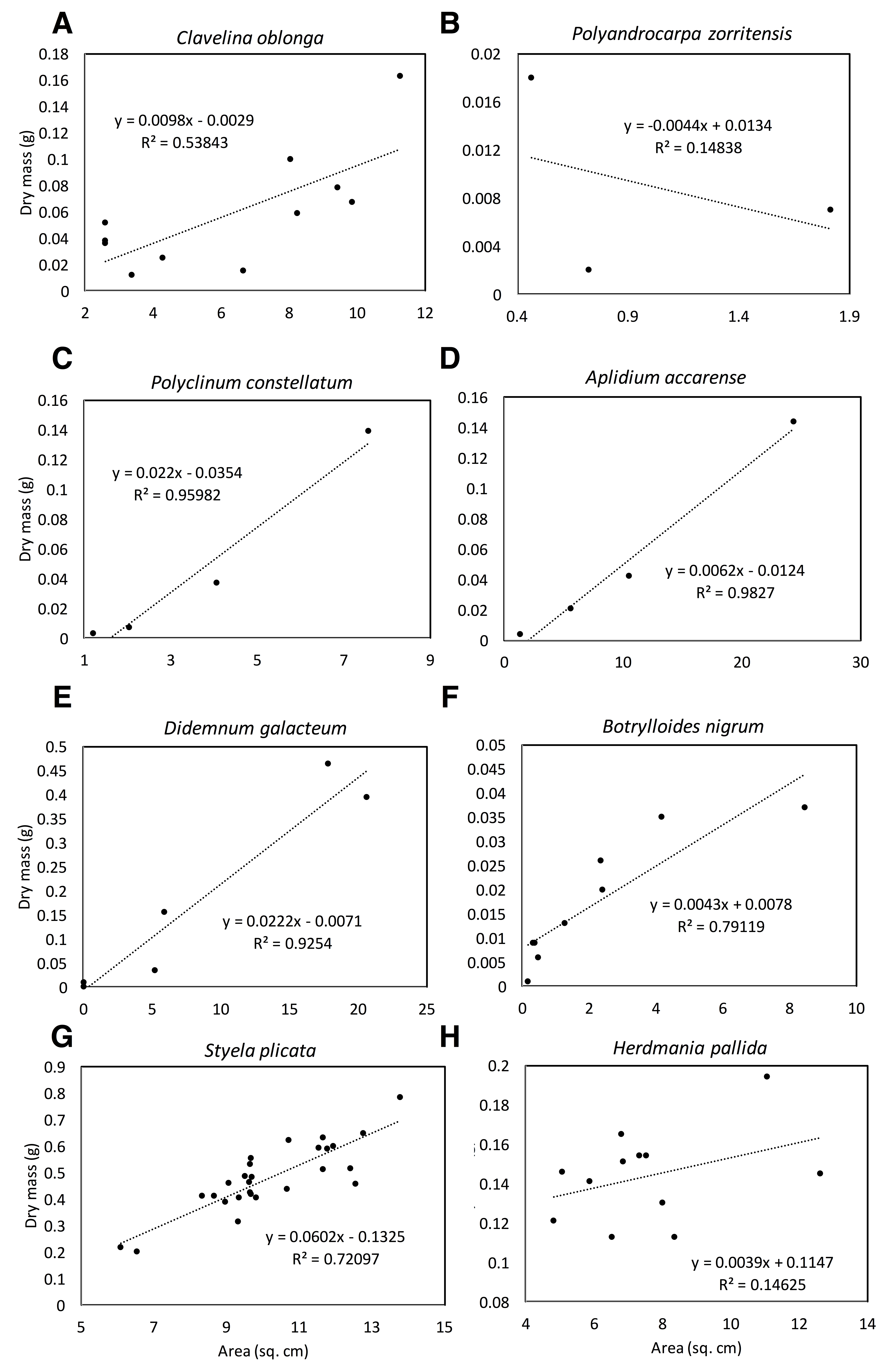
